# Differential roles of beta-cell IP3R and RyR ER Ca^2+^ channels in ER stress-induced alterations of beta-cell Ca^2+^ homeostasis

**DOI:** 10.1101/2022.10.11.511769

**Authors:** Irina X. Zhang, Andrea Hermann, Juan Leon, Anoop Arunagiri, Peter Arvan, Leslie S. Satin

**Affiliations:** Department of Pharmacology and Brehm Diabetes Research Center, University of Michigan Medical School, Ann Arbor, MI 48105; J. Crayton Pruitt Family Department of Biomedical Engineering, University of Florida, Gainesville, FL 32612; Department of Metabolism, Endocrinology & Diabetes, University of Michigan, Ann Arbor, MI 48105

**Keywords:** calcium channels, ryanodine receptor (RyRs), inositol 1,4,5-triphosphate (IP_3_) receptor (IP_3_Rs), endoplasmic reticulum stress (ER stress), beta cells

## Abstract

Pancreatic beta cells maintain glucose homeostasis by secreting pulses of insulin in response to a rise in glucose. Pulsatile secretion occurs due to glucose-induced oscillations in beta-cell cytosolic Ca^2+^. The endoplasmic reticulum (ER) helps regulate beta-cell cytosolic Ca^2+^, and ER stress can lead to ER Ca^2+^ depletion, beta-cell dysfunction and an increased risk of type 2 diabetes. To determine the effects of tunicamycin-induced ER stress on ER inositol 1,4,5-triphosphate receptors (IP3Rs) and ryanodine receptors (RyRs) and their involvement in subsequent Ca^2+^ dysregulation, INS-1 832/13 cells and primary mouse islets were treated with tunicamycin. This increased RyR1 mRNA and potentiated RyR-mediated Ca^2+^ signaling without affecting RyR2 mRNA. TM treatment also enhanced IP3R function, while it decreased IP3R1 and IP3R3 mRNA. Stress reduced ER Ca^2+^, triggered oscillations in cytosolic Ca^2+^ under subthreshold glucose conditions, and increased apoptosis; these changes were prevented by cotreatment with the RyR1 inhibitor dantrolene. In contrast, inhibiting IP3Rs with xestospongin-C failed to suppress the cytosolic Ca^2+^ oscillations due to tunicamycin treatment and did not protect beta cells from tunicamycin-induced apoptosis, although xestospongin-C inclusion prevented ER Ca^2+^ depletion. Taken together, changes in RyR1 function were shown to play a critical role in ER stress induced Ca^2+^ dysfunction and beta-cell apoptosis.

## Introduction

Calcium (Ca^2+^) is an essential cellular signal. In pancreatic beta cells, Ca^2+^ triggers insulin secretion to maintain postprandial blood glucose (Howell, Jones, and Persaud 1994). Ca^2+^ is sequestered within the endoplasmic reticulum (ER), the organelle where the synthesis and folding of secretory proteins occurs, along with lipid synthesis (Berridge 2002; Anelli and Sitia 2008). A well-functioning ER is critical for proper beta-cell function and survival (Fonseca, Gromada, and Urano 2011; Hasnain, Prins, and McGuckin 2016; I. X. Zhang, Raghavan, and Satin 2020; Arunagiri et al. 2019). On the other hand, ER malfunction can potentially lead to type 2 diabetes, and there is evidence that the unfolded protein response (UPR) is activated in islets from type 2 diabetes patients or animal models of diabetes (I. X. Zhang, Raghavan, and Satin 2020; Back and Kaufman 2012).

Decreased ER Ca^2+^ concentration ([Ca^2+^]_ER_) is associated with ER stress and apoptosis, and accompanies many pathologies (I. X. Zhang, Raghavan, and Satin 2020; Sammels et al. 2010; Mekahli et al. 2011). ER Ca^2+^ is regulated by a balance between Ca^2+^ uptake via Sarco/Endoplasmic Reticulum Ca^2+^-ATPases (SERCA pumps) and Ca^2+^ release by inositol 1,4,5-trisphosphate receptors (IP3Rs) and ryanodine receptors (RyRs) (Clapham 2007). Both IP3Rs and RyRs are gated by Ca^2+^, and both trigger Ca^2+^-induced Ca^2+^ release (CICR) under certain conditions (Graves and Hinkle 2003; Dyachok, Tufveson, and Gylfe 2004). There are three known isoforms of IP3Rs: IP3R1, IP3R2 and IP3R3 (16, 27, 28); IP3R1 is the most abundant isoform in beta cells (Ye et al. 2011). RyRs are also encoded by three separate genes, RyR1, RyR2 and RyR3 (Santulli et al. 2017; 2015), with RyR2 being most abundant in beta cells (Yamamoto et al. 2019). Beta-cell IP3Rs have been extensively studied, and their role in GPCR-coupled intracellular Ca^2+^ release is well established (Santulli et al. 2017). In contrast, the role of RyRs has been more limited and controversial, in part because RyR expression in beta cells is very low (Yamamoto et al. 2019).

In this study, we tested whether ER stress differentially regulated RyRs and IP3Rs in beta cells, to better define their respective roles in beta-cell function, such as in ER Ca^2+^ handling and the production of cytosolic Ca^2+^ ([Ca^2+^]_cyto_) oscillations. We used tunicamycin (TM) to experimentally induce ER stress in the rat insulin-secreting INS-1 832/13 beta-cell line, as well as isolated mouse islets. TM activates the UPR by inhibiting N-acetylglucosamine phosphotransferase, leading to protein misfolding in the ER (Yoo et al. 2018). TM treatment triggered [Ca^2+^]_cyto_ oscillations and apoptosis through RyR1-mediated ER Ca^2+^ release. IP3Rs in contrast appeared to play a minor role in these processes. Importantly, while RyR2 has been considered the key isoform regulating ER Ca^2+^ in beta cells (Santulli et al. 2015; Yamamoto et al. 2019), our results emphasize the importance of RyR1 in ER stress. Ultimately, these findings suggest that it might be beneficial to target RyR1 as a potential therapy to alleviate ER stress-mediated beta-cell dysfunction in type 2 diabetes.

## Results

In previous work, we reported that chemically inducing ER stress in beta-cells using TM activated store-operated Ca^2+^ entry (SOCE) and led to the appearance of [Ca^2+^]_cyto_ oscillations in parallel with oscillations in membrane potential (I. X. Zhang et al. 2020). The increase in [Ca^2+^]_cyto_ concomitantly augmented insulin secretion under what would normally be subthreshold glucose conditions, e.g., in medium containing 5 mM glucose (I. X. Zhang et al. 2020). TM induced a reduction in [Ca^2+^]_ER_, which we suggested was likely the proximal trigger for inducing extracellular Ca^2+^ influx via store-operated calcium entry (SOCE). Ca^2+^ can diffuse out of the ER through open IP3Rs, RyRs or possibly the ER translocon (not addressed in this paper; I. X. Zhang, Raghavan, and Satin 2020). In response to ER stress, IP3Rs and RyRs may become dysregulated, resulting in enhanced ER Ca^2+^ efflux. As this possibility was not addressed in our previous work, here we decided to investigate the respective roles of IP3Rs and RyRs in altered beta-cell function under ER stress conditions. To do so, we took advantage of the selective ER Ca^2+^ channel antagonists xestospongin-C (XeC; Gafni et al. 1997) and dantrolene (Dan; Zhao et al. 2001), to block IP3Rs and RyR1, respectively.

We first analyzed the effect of TM on the expression of each of the receptor isoforms. As shown in Figure 1A-C, TM treatment decreased IP3R1 and IP3R3 mRNAs, while the IP3R2 transcript remained unchanged after a 16 h exposure to TM. To assay IP3R function, carbachol was used to activate muscarinic acetylcholine receptors on the beta-cell surface and stimulate IP3Rs by increasing cytosolic IP_3_ (White and McGeown 2002). IP3R-mediated gating of ER Ca^2+^ release is shown in Figure 1D and 1F after normalizing to the initial value. The area under the curve (AUC) of the carbachol responses we observed was increased in TM-treated INS-1 832/13 cells (Figure 1E) and in islets (Figure 1G). As shown in Figure 1H-I, RyR1 mRNA was increased by TM treatment in INS-1 832/13 cells, while the level of RyR2 transcript was unchanged. Caffeine-stimulated ER Ca^2+^ release through RyRs (White and McGeown 2002) was revealed in Figure 1J and 1L after normalizing to the initial value. A 16 h treatment with TM increased the AUC of Ca^2+^ release in both INS-1 832/13 cells (Figure 1K) and islets (Figure 1M). While we wished to measure RyRs and IP3Rs at the protein level, our efforts to do so were hampered by the lack of commercially available, working antibodies for these ER Ca^2+^ channel proteins.

**Figure 1.**
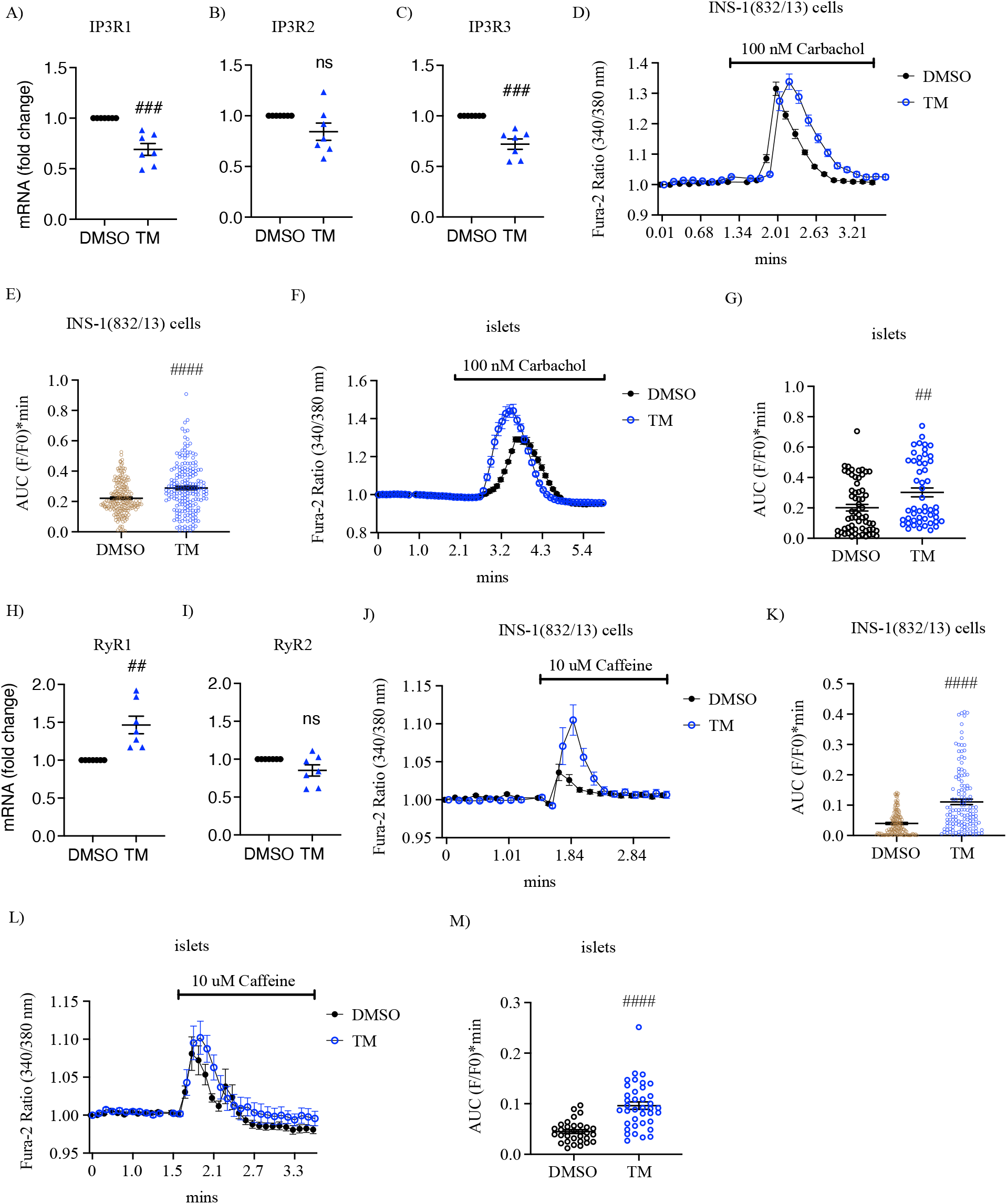
Tunicamycin altered IP3Rs and RyRs expression and function. INS-1 832/13 cells or isolated mouse islets were treated with vehicle control (DMSO) or tunicamycin (TM, 10 μg/ml) for 16 hours in 11 mM glucose INS-1 832/13 culture medium. IP3R isoforms mRNA levels were measured in INS-1 832/13 cells. 1A: IP3R1. 1B: IP3R2. 1C: IP3R3. 1D and 1F: The responses of [Ca^2+^]_cyto_ to the solution containing 11 mM glucose without Ca^2+^ under the indicated conditions. 1E and 1G: Area under the curve (AUC) analysis of Figure 1D and 1F, respectively. RyR isoforms mRNA levels were measured in INS-1 832/13 cells. 1H: RyR1. 1I: RyR2. 1J and 1L: The responses of [Ca^2+^]_cyto_ to the solution containing 11 mM glucose without Ca^2+^ under the indicated conditions. [Ca^2+^]_cyto_ was normalized to its initial Fura-2 ratio. 1K and 1M: Area under the curve (AUC) analysis of 1J and 1L, respectively. All values shown are means ± SEM, #, p< 0.05, ##, p< 0.01, ###, p< 0.005, ####, p< 0.0001; n= 4-7 times repeated per condition, by student’s t-test.

To test whether blocking IP3Rs or RyRs prevented the TM-mediated depletion of ER Ca^2+^ (I. X. Zhang et al. 2020), the ER Ca^2+^ probe D4ER was transiently expressed in islets with adenovirus. Islets were then treated with either vehicle control (DMSO), TM, XeC, Dan, TM+XeC or TM+Dan for 16 h, and [Ca^2+^]_ER_ was measured using a recording solution containing 5 mM glucose. Representative traces show [Ca^2+^]_ER_ (Y-axis, in arbitrary units) and the effect of the reversible SERCA blocker cyclopiazonic acid (CPA, 50 µM) depleted ER Ca^2+^, as expected (Laursen et al. 2009) (Figure 2A). Figure 2B-C shows steady-state [Ca^2+^]_ER_ measured in 5 mM glucose in the absence of CPA. Exposing islets to TM for 16 h significantly reduced steady-state [Ca^2+^]_ER_ compared to controls, and the presence of either XeC or Dan with TM reduced ER Ca^2+^ loss. Interestingly, Dan inclusion with TM slightly increased [Ca^2+^]_ER_, but not in controls perhaps due to ER Ca^2+^ leak via RyRs was lower in controls, so that Dan had less effect than it did in

**Figure 2.**
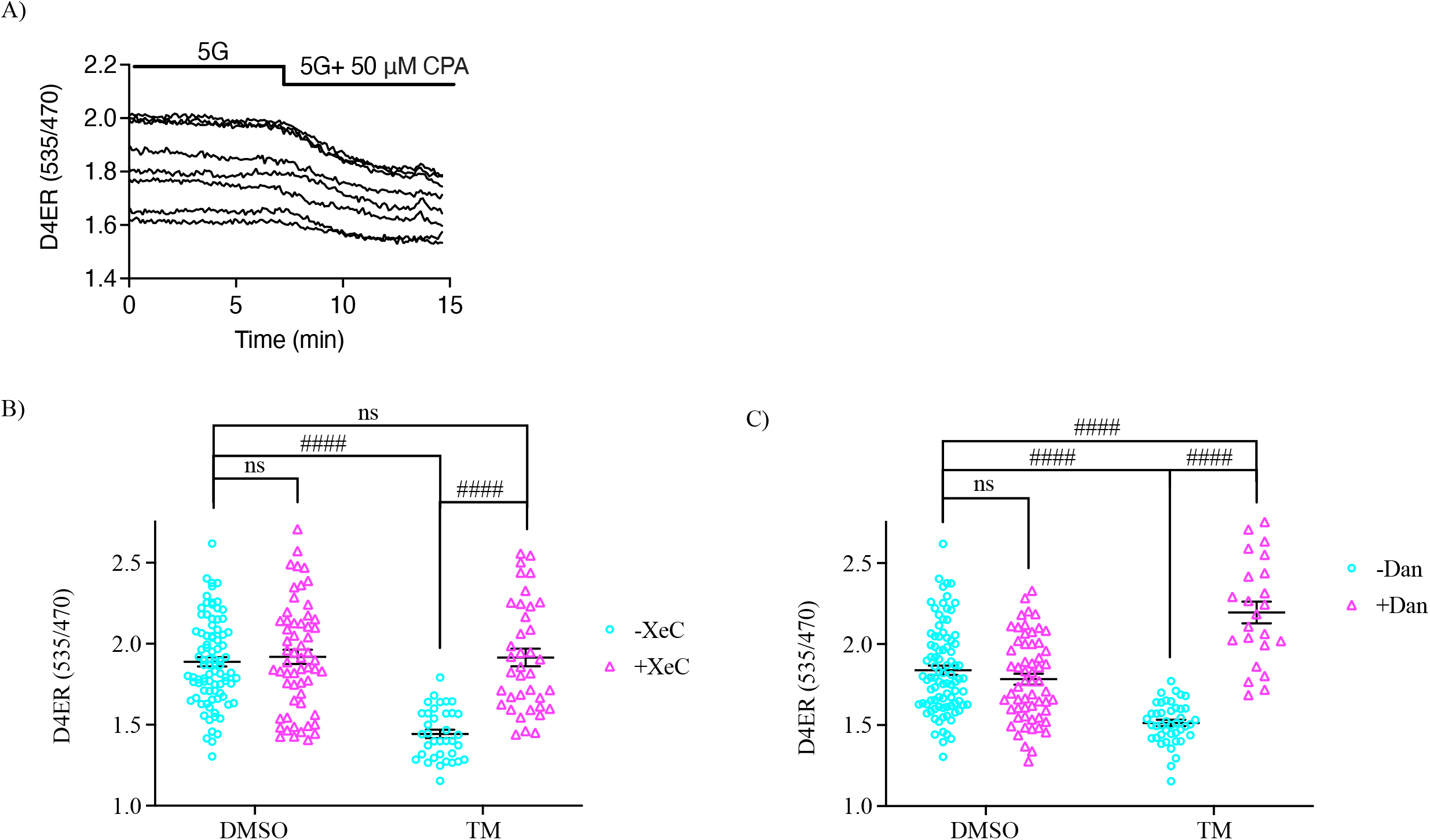
Xestospongin C or dantrolene restored basal [Ca^2+^]_ER_ in tunicamycin-treated islets. Mouse pancreatic islets were infected with an adenovirus expressing a beta-cell directed D4ER probe for three hours, followed by a 48-hour recovery period. Islets were then treated with vehicle control (DMSO), tunicamycin (TM, 10 μg/ml), xestosponginc C (XeC, 1 µM), dantrolene (Dan, 10 µM), TM+ XeC or TM+ dantrolene, for 16 hours in 11 mM glucose islet culture medium. 2A: Representative basal [Ca^2+^]_ER_ traces for control obtained in 5 mM glucose solution before and after cyclopiazonic acid (CPA, 50 μM) application. 2B and 2C: Basal [Ca^2+^]_ER_ for indicated conditions in 5 mM glucose solution. Each data point shown was a D4ER ratio obtained for one selected region of interest, a single cell or small group of cells. 2B: Row Factor F(1, 214)= 30.21, p< 0.0001, Column Factor F(1, 214)= 37.83, p< 0.0001, Interaction F(1, 214)= 29.13, p< 0.0001. 2C: Row Factor F(1, 211)= 1.242, p= 0.2663, Column Factor F(1, 211)= 65.89, p< 0.0001, Interaction F(1, 211)= 90.98, p< 0.0001. All values shown are means ± SEM. ####, p< 0.0001; ns= not significant; n= at least 3 mice, by two-way ANOVA with post hoc multiple comparison by Tukey’s procedure.

TM-treated islets (Figure 2C).

To determine whether IP3Rs and/or RyRs play a role in the production of the oscillations seen in subthreshold glucose in stressed beta cells, islets were exposed to a vehicle control (DMSO), TM, XeC, Dan, TM+XeC or TM+Dan for 16 h before recording [Ca^2+^]_cyto_ in a solution containing 5 mM glucose. Mouse islets cultured overnight in media containing 11 mM glucose do not typically exhibit oscillations in [Ca^2+^]_cyto_ when acutely exposed to subthreshold glucose levels (i.e., glucose concentrations < 7 mM) (I. X. Zhang et al. 2020; Satin et al. 2015; Glynn et al. 2016). As expected, control islets displayed little or no oscillatory [Ca^2+^]_cyto_ (Figure 3). In contrast, ~80% of TM-treated islets exhibited Ca^2+^ oscillations in 5 mM glucose (Figure 3) due to the activation of SOCE, as demonstrated in our previous study, even though these channels are not normally involved in the production of glucose-induced islet oscillations under physiological conditions (I. X. Zhang et al. 2020; Bertram, Satin, and Sherman 2018). Including XeC with TM had no effect on the production of Ca^2+^ oscillations in stressed islets (Figure 3A and 3C); however, TM-triggered oscillations were suppressed when Dan was present (< 5% oscillating islets) (Figure 3B and 3D), suggesting RyRs but not IP_3_Rs were involved in their genesis. In Dan or XeC-treated islets, as for DMSO, little or no oscillatory Ca^2+^ activity was observed in the absence of TM (Figure 3).

**Figure 3.**
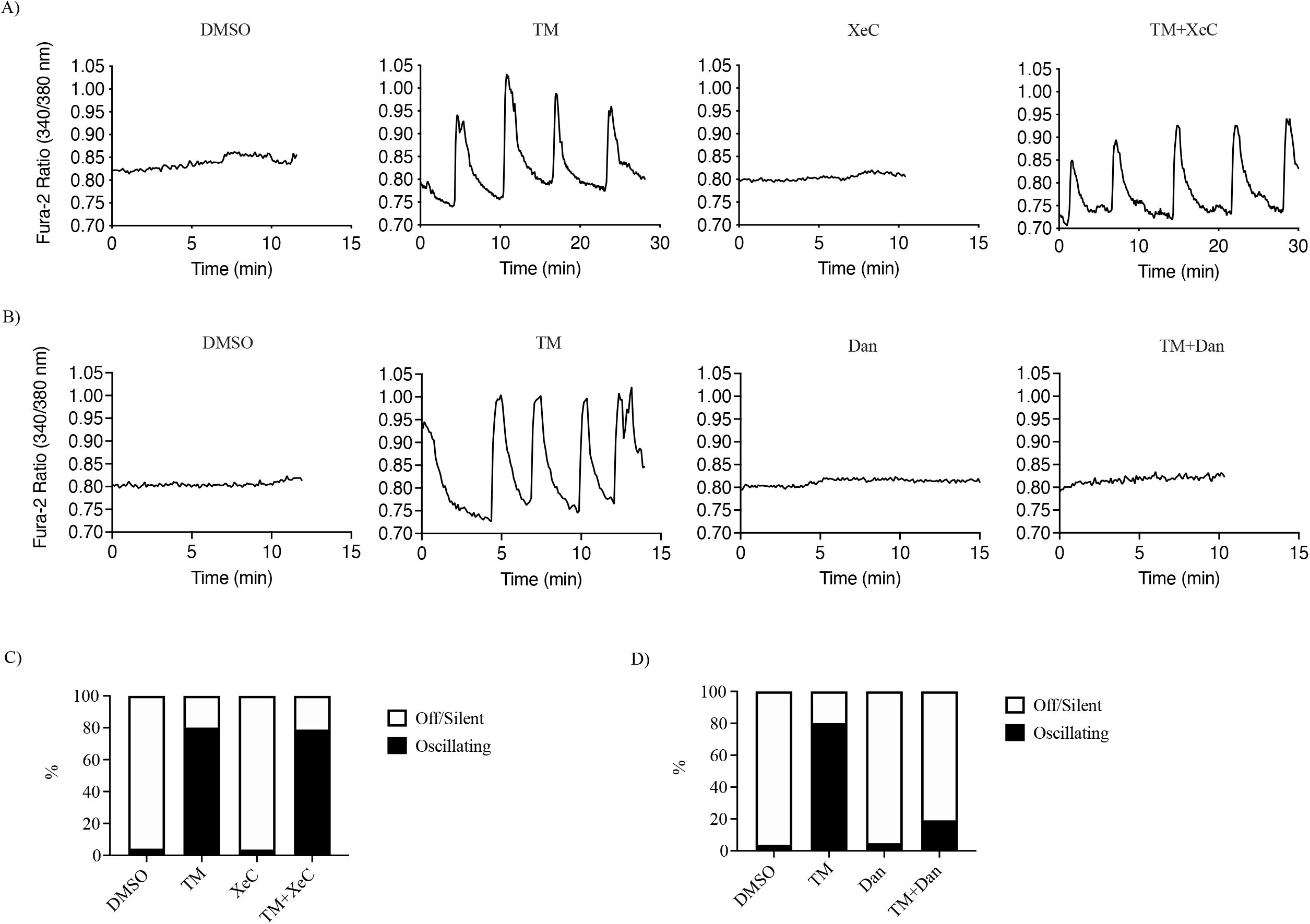
Differential effects of xestospongin C and dantrolene on [Ca^2+^]_cyto_ oscillations under subthreshold glucose conditions. Isolated pancreatic mouse islets were treated with vehicle control (DMSO), tunicamycin (TM, 10 μg/ml), 3A and 3C: xestospongin C (XeC, 1 µM) or TM+XeC; 3B and 3D: dantrolene (Dan, 10 µM) or TM+Dan for 16 hours in 11 mM glucose islet culture medium. 3A and 3B: The responses of [Ca^2+^]_cyto_ to the solution containing 5 mM glucose under the indicated conditions. 3C and 3D: Percentage of oscillating islets; n= at least 3 mice.

To confirm that RyR1 played a role in the production of TM-triggered [Ca^2+^]_cyto_ oscillations in 5 mM glucose, we transfected RyR1 siRNA in INS-1 832/13 cells, resulting in a 50% decrease of RyR1 mRNA (Figure 4A). RyR function was assessed by acutely exposing the transfected cells to caffeine in [Ca^2+^]_cyto_ imaging buffer in the absence of Ca^2+^ to isolate ER Ca^2+^ efflux from possible Ca^2+^ influx (Figure 4B). The AUC measurements of the resulting Ca^2+^ recordings were decreased in the siRyR1-cells compared to cells transfected with negative control siRNA (siCon) (Figure 4C). Decreasing RyR1 expression resulted in an increase in RyR2 mRNA levels (Figure 4D), likely as a compensation. Transfected cells were recorded using imaging buffer containing 5 mM glucose (Figure 4E). After a 16 h of TM treatment, the percentage of cells displaying [Ca^2+^]_cyto_ oscillations was decreased in RyR1-knockdown cells (~10%) compared to siCon-cells (~40%) (Figure 4F). RyR1-knockdown did not affect the Ca^2+^ activity of unstressed control cells (Figure 4E-4F). Following RyR1-knockdown, IP3R1 mRNA and IP3R function were both found to be elevated (Figures 4G-I). INS-1 832/13 cells transfected with either negative control or siRNA-RyR1 were exposed to carbachol in 0 Ca^2+^ imaging buffer to test IP_3_R function; AUCs were greater in RyR1-knockdown cells compared to siCon-treated cells, presumably to compensate for RyR1 suppression (Figure 4H-I).

**Figure 4.**
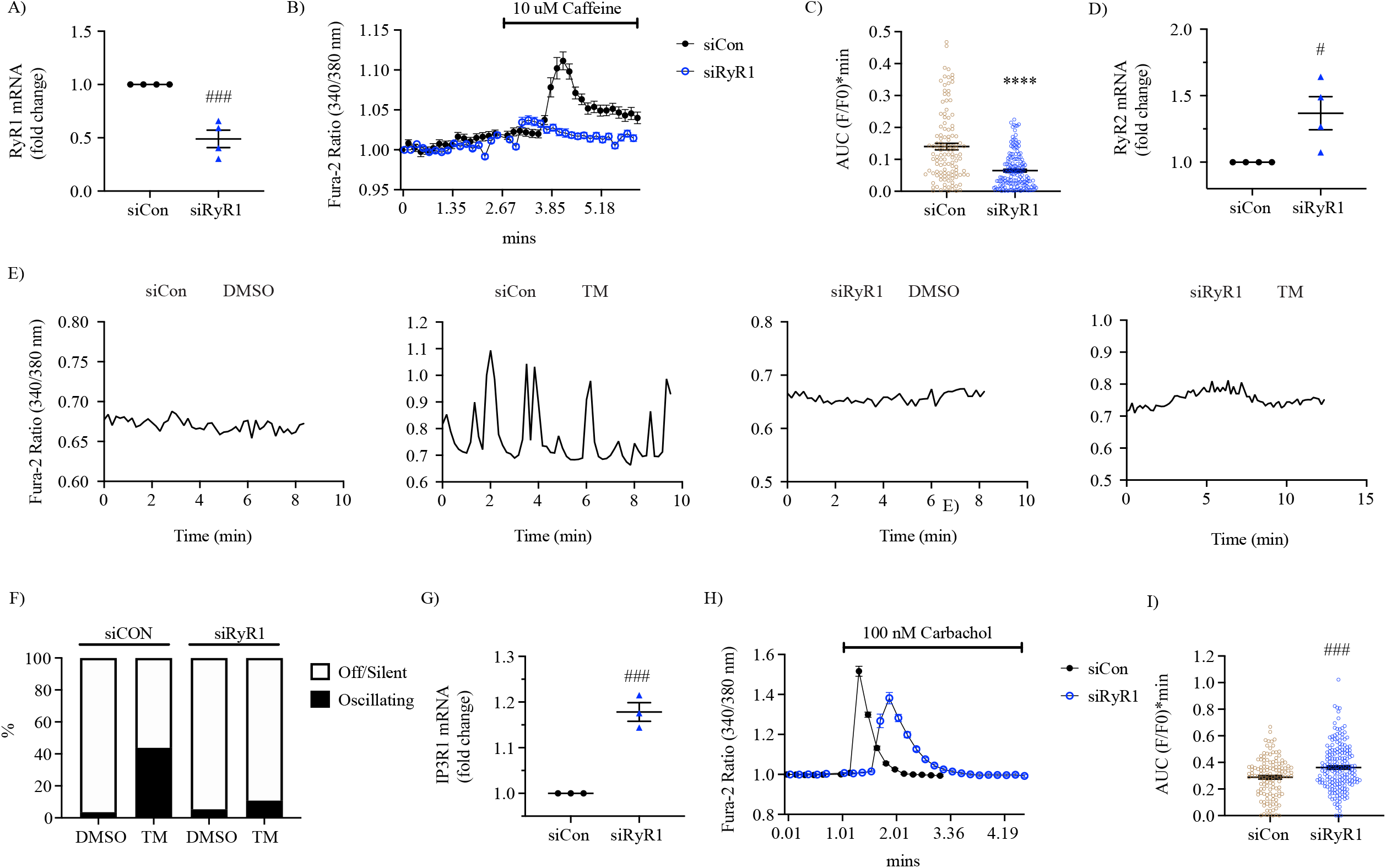
RyR1-knockdown inhibited tunicamycin-triggered [Ca^2+^]_cyto_ oscillations. 4A: RyR1-knockdown in INS-1 832/13 cells was assessed by qPCR 64 hours after siRNA transfection. 4B: The responses of [Ca^2+^]_cyto_ to 10 µM caffeine in the solution containing 11 mM glucose without Ca^2+^ under the indicated conditions. [Ca^2+^]_cyto_ was normalized to its initial Fura-2 ratio. 4C: Area under the curve (AUC) analysis of 4B. 4D: Expression level of RyR2 mRNA in INS-1 832/13 cells after transfection with RyR1 siRNA. INS-1 832/13 cells were treated with vehicle control (DMSO) or tunicamycin (TM, 10 µg/ml) for 16 hours after transfecting with RyR1 siRNA or negative control siRNA for 48 hours. 4E: The responses of [Ca^2+^]_cyto_ to solution containing 5 mM glucose. 4F: Percentage of oscillating INS-1 832/13 cells. 4G: Expression level of IP3R1 mRNA in INS-1 832/13 cells after transfection with RyR1 siRNA. 4H: The responses of [Ca^2+^]_cyto_ to 100 nM carbachol in the solution containing 11 mM glucose without Ca^2+^ under the indicated conditions. [Ca^2+^]_cyto_ was normalized to its initial Fura-2 ratio. 4I: Area under the curve (AUC) analysis of 4H. All values shown are means ± SEM. ##, p< 0.01; ####, p< 0.0001; n= at least 3 times repeated per condition.

RyR2, rather than RyR1, is the predominant RyR isoform of beta-cell in terms of abundance (Yamamoto et al. 2019). To study whether RyR2 is also involved in TM-induced [Ca^2+^]_cyto_ oscillations in 5 mM glucose, we silenced RyR2 using siRNA in INS-1 832/13 cells and achieved a ~50% reduction in RyR2 mRNA (Figure 5A) and a significant reduction of RyR function. RyR2 knockdown cells did not respond to caffeine, and the AUC of the Ca^2+^ recordings was miniscule (Figure 5B-C). RyR1 mRNA remained unchanged after RyR2 knockdown (Figure 5D). The cytosolic Ca^2+^ levels of the transfected cells were recorded using imaging buffer containing 5 mM glucose (Figure 5E). The percentage of cells displaying [Ca^2+^]_cyto_ oscillations in response to a 16 h of TM treatment in this case was unaffected by silencing RyR2 (Figure 5F). Statistically, IP3R1 mRNA levels did not change significantly following RyR2 knockdown (Figure 5G), while IP3R function was increased, as indicated by the greater AUC observed in RyR2-knockdown cells compared to control (Figure 5H-I).

**Figure 5.**
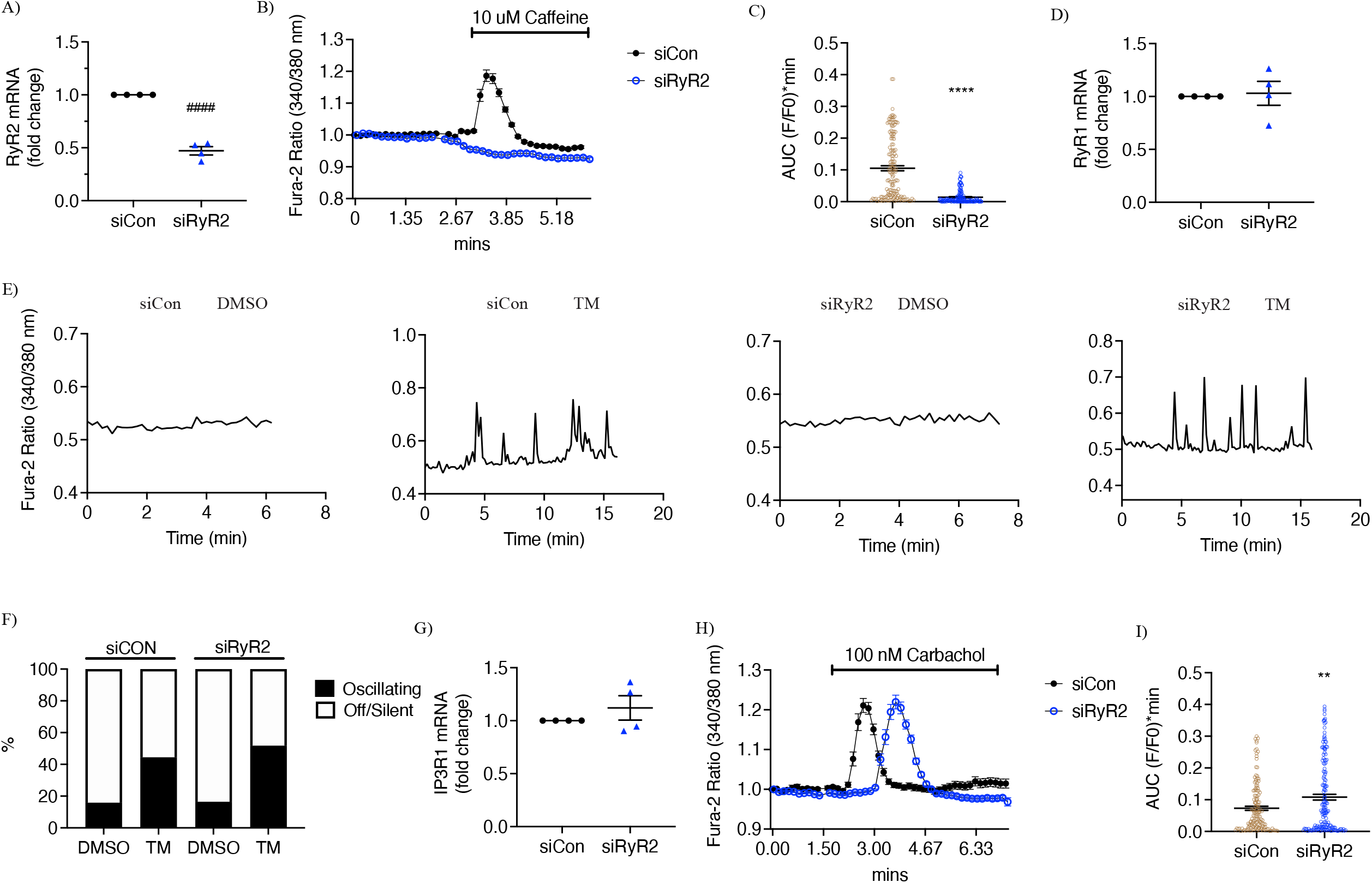
RyR2-knockdown did not affect tunicamycin-triggered [Ca^2+^]_cyto_ oscillations. 5A: RyR2-knockdown in INS-1 832/13 cells was assessed by qPCR 64 hours after siRNA transfection. 5B: The responses of [Ca^2+^]_cyto_ to 10 µM caffeine in the solution containing 11 mM glucose without Ca^2+^ under the indicated conditions. [Ca^2+^]_cyto_ was normalized to its initial Fura-2 ratio. 5C: Area under the curve (AUC) analysis of 5B. 5D: Expression level of RyR1 mRNA in INS-1 832/13 cells after transfection with RyR2 siRNA. INS-1 832/13 cells were treated with vehicle control (DMSO) or tunicamycin (TM, 10 µg/ml) for 16 hours after transfecting with RyR2 siRNA or negative control siRNA for 48 hours. 5E: The responses of [Ca^2+^]_cyto_ to solution containing 5 mM glucose. 5F: Percentage of oscillating INS-1 832/13 cells. 5G: Expression level of IP3R1 mRNA in INS-1 832/13 cells after transfection with RyR2 siRNA. 5H: The responses of [Ca^2+^]_cyto_ to 100 nM carbachol in the solution containing 11 mM glucose without Ca^2+^ under the indicated conditions. [Ca^2+^]_cyto_ was normalized to its initial Fura-2 ratio. 5I: Area under the curve (AUC) analysis of 5H. All values shown are means ± SEM. #, p< 0.05, ###, p< 0.0005; ####, p< 0.0001; n= at least 3 times repeated per condition.

Activation of the UPR occurs in response to TM in many cell types, including beta-cells (I. X. Zhang et al. 2020; Yamamoto et al. 2019; Collett et al. 2018). We previously showed that increases in several of the canonical markers of the ER stress response, such as spliced XBP1, CHOP and BiP, occurred in TM-treated INS-1 832/13 cells and mouse islets (I. X. Zhang et al. 2020). To better define the respective roles of IP3Rs and RyRs in TM-induced ER stress in beta-cell, INS-1 832/13 cells were treated with TM (10 µg/ml) or vehicle control (DMSO) with or without XeC (1µM) or Dan (10 µM) for 6 h. Total mRNA was then extracted and quantified, as we had previously observed that spliced XBP1 reached its peak at this time (I. X. Zhang et al. 2020). However, neither XeC nor Dan suppressed UPR activation, as indicated by an increased ratio of spliced XBP1/total XBP1 (Figure 6A and 6D), ATF4 (Figure 6B and 6E) or CHOP (Figure 6C and 6F), which we observed when these inhibitors were included with TM. Thus, blocking IP3Rs or RyR1 failed to prevent UPR activation in TM-treated beta-cells.

**Figure 6.**
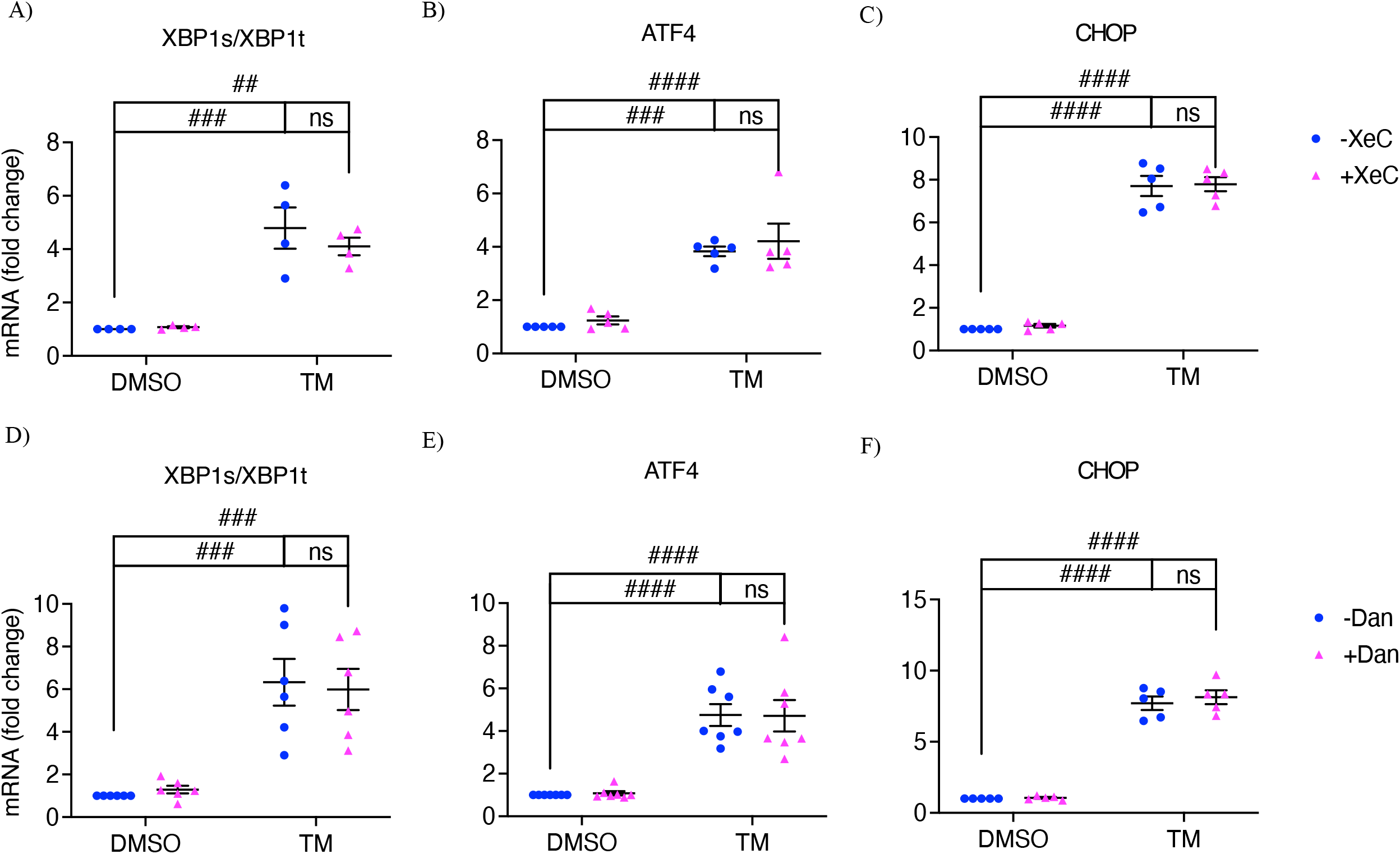
Xestospongin C or dantrolene did not attenuate UPR activation. INS-1 832/13 cells were treated with vehicle control (DMSO), tunicamycin (TM, 10 µg/ml), 6A-6C: xestospongin C (XeC, 1 µM) or TM+XeC; 6D-6F: dantrolene (Dan, 10 µM) or TM+Dan for 6 hours in 11 mM glucose INS-1 832/13 culture medium. Various ER stress markers were measured. 6A and 6D: spliced XBP1/total XBP1 ratio. 6B and 6E: ATF4. 6C and 6F: CHOP. 6A: Row Factor F(1, 12)= 65.48, p< 0.0001, Column Factor F(1, 12)= 0.5282, p= 0.4813, Interaction F(1, 12)= 0.8167, p= 0.3839. 6B: Row Factor F(1, 16)= 68.82, p< 0.0001, Column Factor F(1, 16)= 0.7796, p= 0.3903, Interaction F(1, 16)= 0.03948, p= 0.8450. 6C: Row Factor F(1, 16)= 530.0, p< 0.0001, Column Factor F(1, 16)= 0.1767, p= 0.6798, Interaction F(1, 16)= 0.01755, p= 0.8963. 6D: Row Factor F(1, 20)= 46.47, p< 0.0001, Column Factor F(1, 20)= 0.0009452, p= 0.9758, Interaction F(1, 20)= 0.1819, p= 0.6743. 6E: Row Factor F(1, 24)= 66.73, p< 0.0001, Column Factor F(1, 24)= 0.002392, p= 0.9614, Interaction F(1, 24)= 0.01586, p= 0.9008. 6F: Row Factor F(1, 16)= 412.3, p< 0.0001, Column Factor F(1, 16)= 0.5069, p= 0.4868, Interaction F(1, 16)= 0.3036, p= 0.5892. All values shown are means ± SEM. ##, p< 0.01; ###, p< 0.005; ####, p< 0.0001; ns= not significant; n= at least 4 times repeated per condition, by two-way ANOVA with post hoc multiple comparison by Tukey’s procedure.

To confirm that blocking RyRs does not affect TM-triggered UPR activation, we assessed ATF4 and CHOP expression in RyR1, RyR2 or RyR1+2 knockdown cells, and observed that neither ATF4 or CHOP upregulation was reversed by silencing these isoforms individually or collectively (Figure 7A-B). To further confirm this, we also examined the effect of blocking RyRs on UPR activation using ryanodine (Ry), a classic non-specific RyRs blocker [at 100 µM, Ry is a RyR blocker, while at nanomolar concentrations, Ry is a RyR activator (Van Petegem 2012)]. As shown in Figure 7C-D, when INS-1 832/13 cells were treated with TM for 6 h, neither the ratio of spliced XBP1/total XBP1 or CHOP mRNA were reduced by the inclusion of 100 µM of Ry, which was similar what we observed upon RyR1+2 knockdown (Figure 7A-B).

**Figure 7.**
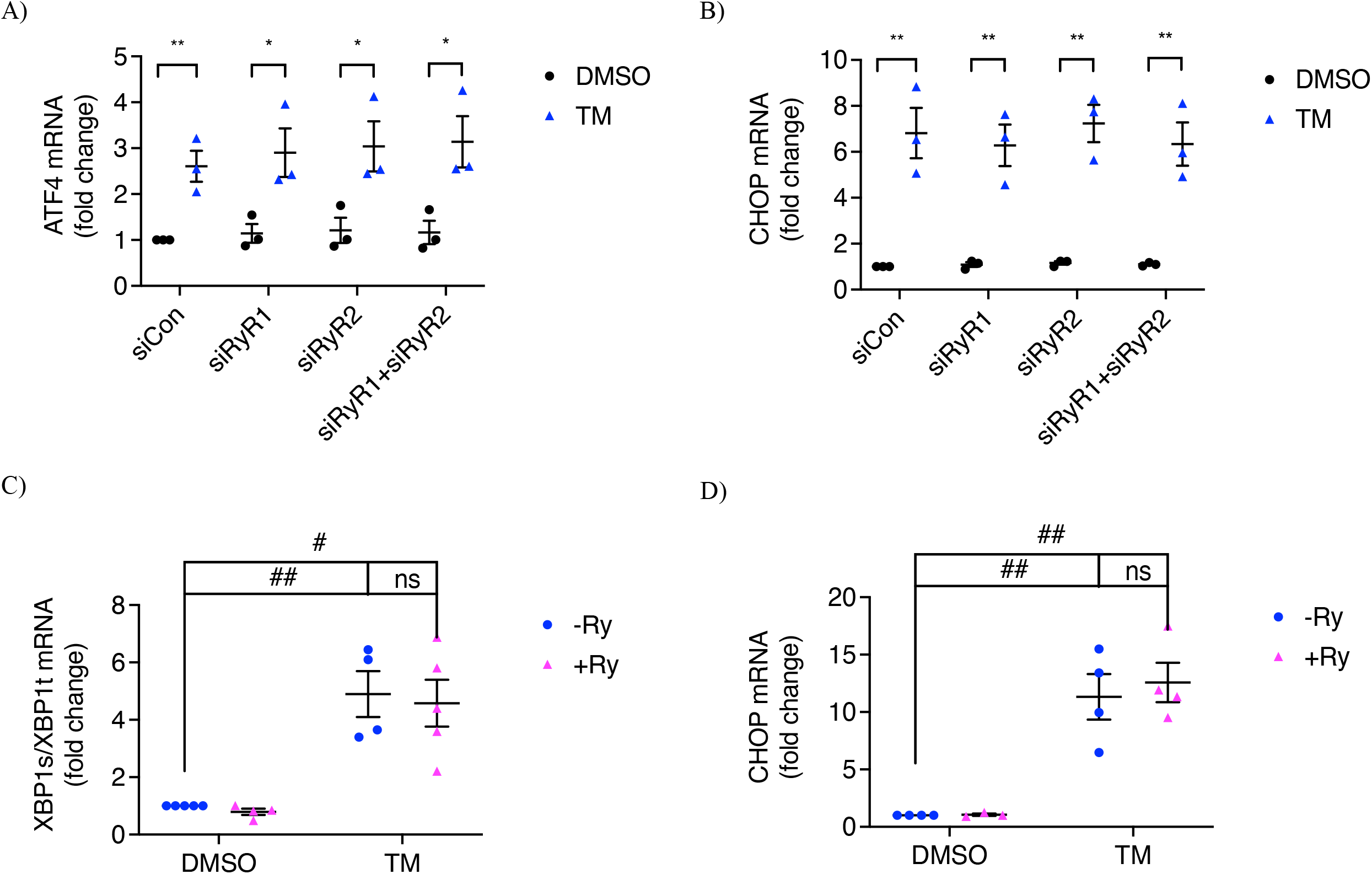
Inhibiting ryanodine receptors did not affect TM-triggered UPR activation. INS-1 832/13 cells were transfected with siRNA for control, RyR1, RyR2 or RyR1+RyR2 for 64 hours and treated with vehicle control (DMSO) or TM (10 µg/ml) for 6 hours in 11 mM glucose INS-1 832/13 culture medium. 7A: ATF4 and 7B: CHOP were measured. INS-1 832/13 cells were treated with vehicle control (DMSO), ryanodine (RyR, 100 µM), tunicamycin (TM, 10µg/ml) or TM+Ry for 6 hours in 11 mM glucose INS-1 832/13 culture medium. 7C: spliced XBP1/total XBP1 ratio; 7D: CHOP were measured. 7C: Row Factor F(1, 14)= 43.32, p< 0.0001, Column Factor F(1, 14)= 0.2020, p= 0.6600, Interaction F(1, 14)= 0.00882, p= 0.9265. 7D: Row Factor F(1, 11)= 58.68, p< 0.0001, Column Factor F(1, 11)= 0.2081, p= 0.6571, Interaction F(1, 11)= 0.1730, p= 0.6855. All values shown are means ± SEM. #, p< 0.05, ##, p< 0.01; ###, p< 0.005; ####, p< 0.0001; ns= not significant; n= at least 3 times repeated per condition, 7A-7B: by student’s t-test; 7C-7D: by two-way ANOVA with post hoc multiple comparison by Tukey’s procedure.

We also treated islets with a much lower dose of TM (300 nM) for 24 h and verified if such low dose of TM could trigger UPR activation. We observed that 90% of TM-treated islets exhibited subthreshold Ca^2+^ oscillations, and the inclusion of Dan lowered the percentage to 10% (Figure S1A-B). In addition, we saw increased ratio of spliced XBP1/total XBP1, ATF4 and CHOP mRNA after a 6 h of treatment that were not affected by Dan (Figure S1C-E). Although there seemed to show a trend of Ry suppressing TM-induced upregulation of XBP1s/XBP1t ratio and CHOP mRNA, but the difference was not statistically significant (Figure S1F-G).

We previously reported that both thapsigargin (TG, 200 nM) and high glucose (HG, 25 mM), applied separately for 16 h, induced subthreshold [Ca^2+^]_cyto_ oscillations in mouse islets (I. X. Zhang et al. 2020). Here we tested whether XeC or Dan affected the oscillations. As shown in Figure S2C, XeC slightly decreased the percentage of islets exhibiting oscillations in response to TG treatment (from ~85% to ~65%), while Dan reduced the percentage to <40%. Similarly, XeC reduced the percentage of islets exhibiting oscillations seen following HG treatment from ~70% to ~40%, while Dan decreased it further to ~20% (Figure S2D).

Programmed cell death or apoptosis has been shown to occur in beta cells in response to prolonged ER stress (I. X. Zhang, Raghavan, and Satin 2020; I. X. Zhang et al. 2020; Oslowski and Urano 2011; Sano and Reed 2013). We examined how XeC or Dan affected the percentage of cells in the sub-G1 phase of the cell cycle, a measure of cell entry into late-stage apoptosis (Riccardi and Nicoletti 2006). As expected at 24 h, TM significantly increased the percentage of INS-1 832/13 cells undergoing apoptosis compared to DMSO- or Dan/XeC-treated controls (Figure 8). XeC+TM was without effect, but Dan+TM considerably reduced apoptosis to near control levels, suggesting a potential linkage between RyR1 and TM-triggered beta-cell apoptosis.

**Figure 8.**
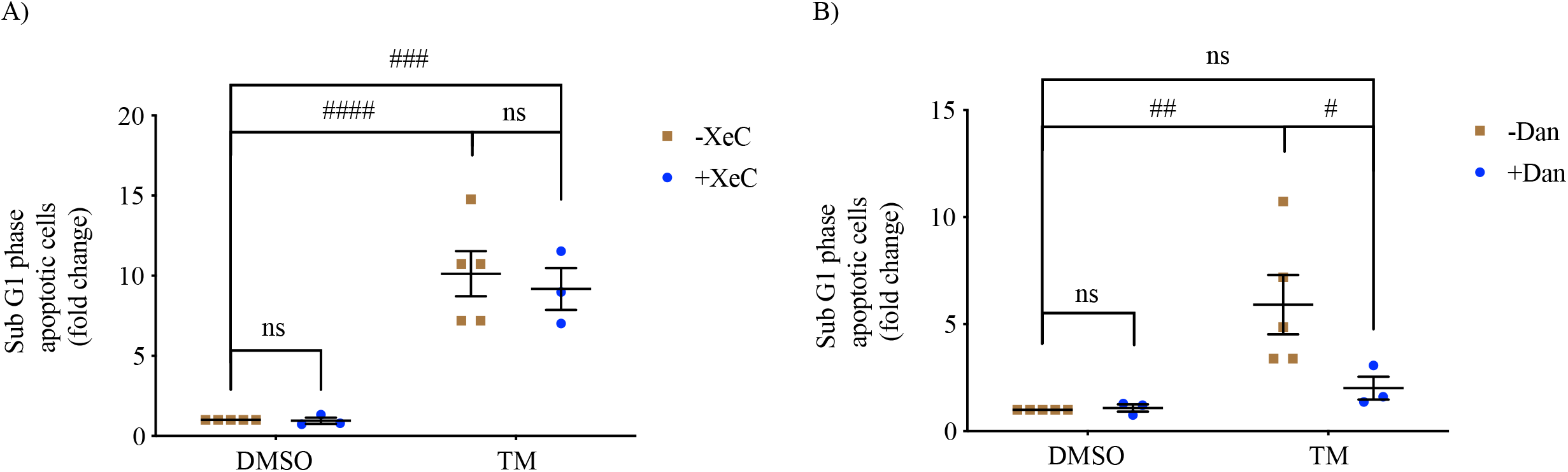
Differential effects of xestospongin C and dantrolene on beta-cell apoptosis. INS-1 832/13 cells were treated with vehicle control (DMSO) or tunicamycin (TM, 10 μg/ml), 8A: xestospongin C (XeC, 1 µM), or TM+XeC; 8B: dantrolene (Dan, 10 µM) or TM+Dan for 24 hours in 11 mM glucose INS-1 832/13 culture medium. Late-stage apoptotic INS-1 832/13 cells is shown using the sub-G1 assay measured by flow cytometry. Fold change was derived by comparing to DMSO group. 8A: Row Factor F(1, 8)= 50.25, p= 0.0001, Column Factor F(1, 8)= 0.4793, p= 0.5083, Interaction F(1, 8)= 0.4267, p= 0.5319. 8B: Row Factor F(1, 8)= 26.26, p= 0.0009, Column Factor F(1, 8)= 5.753, p= 0.0433, Interaction F(1, 8)= 6.879, p= 0.0305. All values shown are means ± SEM. #, p< 0.05, ##, p< 0.01, ###, p< 0.005; ####, p< 0.0001, ns= not significant; n= at least 3 times repeated per condition, by two-way ANOVA with post hoc multiple comparison by Tukey’s procedure.

To test the possible relevance of these results to diabetes, we extended our study to islets isolated from *db/db* mice, an animal model of type 2 diabetes. These mice harbor a mutation in the leptin receptor and have been shown to exhibit a left-shift in their glucose sensitivity over time (Corbin et al. 2016). The mice rapidly gained weight (34.73 ± 2.885g SEM) compared to their heterozygous controls (19.30 ± 2.352g SEM), and developed hyperglycemia (312.3 ± 12.44 mg/dl SEM, n=3), while *db/+* mice maintained normal blood sugar (166 ± 39.93 mg/dl SEM, n=3). After isolation, islets from the mice were treated with control DMSO or Dan for 16 h before [Ca^2+^]_cyto_ measurements were carried out. As shown in Figure 9A, islets from *db/+* mice treated with vehicle lacked oscillations in 5 mM glucose, and as expected, exhibited normal oscillations in response to 11 mM glucose; these responses were unaffected by pretreatment with Dan. In contrast, *db/db* mice exhibited oscillations in 5 mM glucose and plateaus in 11 mM glucose. In these islets, the inclusion of Dan abolished the oscillations and decreased the percentage of oscillating islets observed from 70% to 20% in 5 mM glucose, without affecting the plateaus seen in 11 mM glucose (Figure 9B).

**Figure 9.**
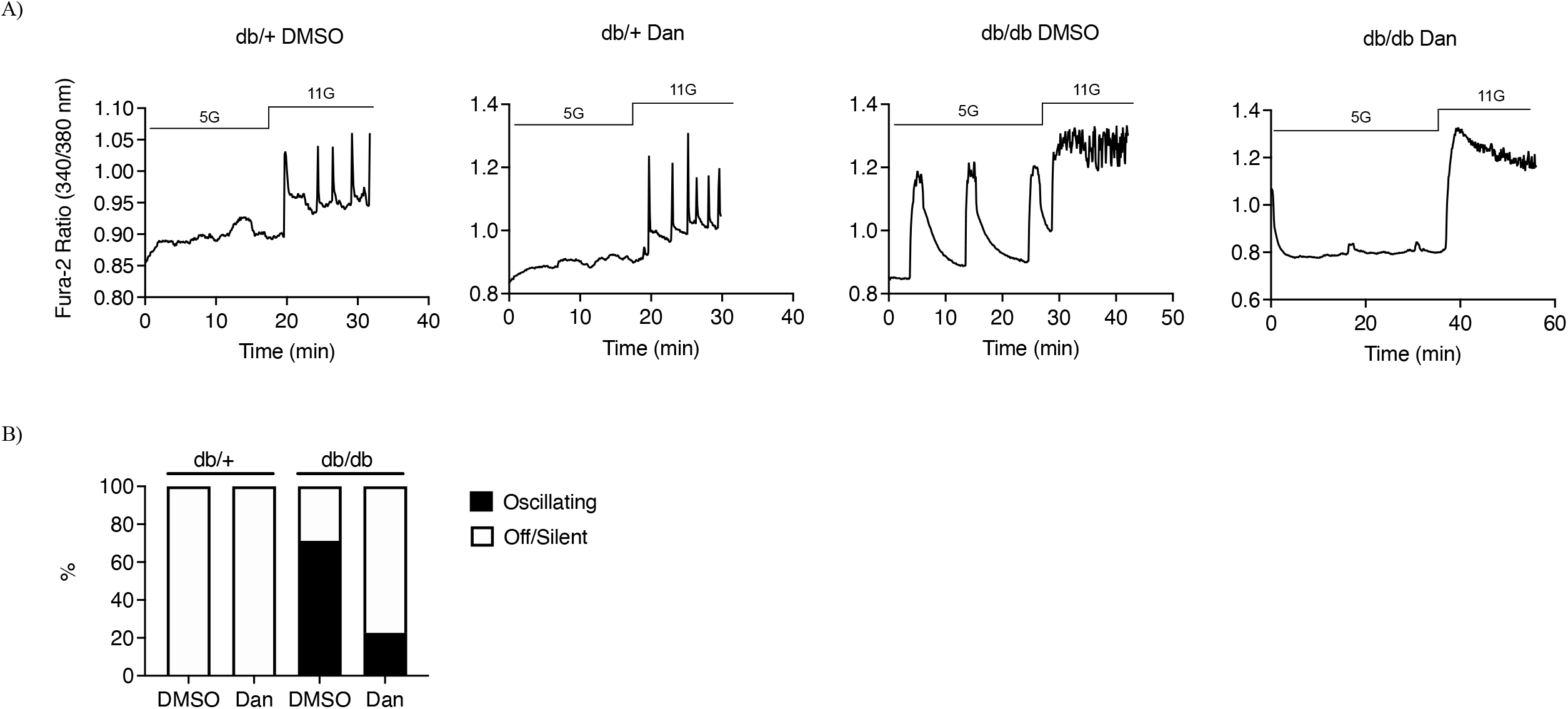
Dantrolene suppressed subthreshold [Ca^2+^]_cyto_ oscillation in islets from *db/db* mice. Pancreatic islets were isolated from female *db/+* and *db/db* mice (5-7 weeks old) and treated with control DMSO or dantrolene (Dan, 10 µM) for 16 hours in 11 mM glucose islet culture medium. 9A: The responses of [Ca^2+^]_cyto_ to the solution containing 5 mM and 11 mM glucose under the indicated conditions. 9B: Percentage of oscillating islets; n= 3 mice.

## Discussion

We previously reported that [Ca^2+^]_cyto_ oscillations and increased insulin secretion observed in ER-stressed beta cells in subthreshold glucose are triggered by ER Ca^2+^ depletion and SOCE activation (I. X. Zhang et al. 2020). However, how ER stress depletes ER Ca^2+^ has remained unclear. Therefore, in the present study, we attempted to differentiate possible roles for RyRs and IP3Rs in ER Ca^2+^ depletion and its downstream effects on beta-cell Ca^2+^ homeostasis. Taken together, the data support the hypothesis that SOCE activation occurs secondary to RyR1-mediated ER Ca^2+^ depletion.

While blocking IP3Rs with XeC prevented a loss in [Ca^2+^]_ER_ in TM-treated islets (Figure 2B); XeC failed to inhibit TM-induced subthreshold [Ca^2+^]_cyto_ oscillations (Figure 3C) or beta-cell apoptosis (Figure 8A). On the other hand, blocking RyR1 with Dan suppressed TM-induced subthreshold [Ca^2+^]_cyto_ oscillations (Figure 3D) and beta-cell apoptosis (Figure 8B) in addition to preventing the reduction in [Ca^2+^]_ER_ induced by TM (Figure 2C). Similar results were reported in a study of Dan as a potential treatment for Wolfram syndrome, a rare autosomal recessive disorder that is associated with childhood onset diabetes mellitus as well as sensory and neurological deficits (Lu et al. 2014). The causative genes for Wolfram syndrome are *WFS1* and *WFS2*, which encode ER-resident proteins. Loss of *WFS1* or *WFS2* has been shown to cause ER stress and decrease [Ca^2+^]_ER_ in neurons (Li et al. 2020). Lu et al. found that knocking down WFS1 in INS-1 832/13 cells or NSC34 cells increased [Ca^2+^]_cyto_ and induced cell death, and importantly, these changes were suppressed by Dan (10 µM, 24 h). Moreover, Dan inclusion (10 µM, 48 h) also inhibited thapsigargin-induced cell death in neural progenitor cells derived from the iPSCs of a Wolfram syndrome patient (Lu et al. 2014).

In addition to inducing chemical ER stress with agents like TM, we also examined the effect of Dan on [Ca^2+^]_cyto_ in islets exposed to high glucose (25 mM, 16 h) to mimic glucose toxicity, and found that Dan suppressed high glucose-triggered subthreshold [Ca^2+^]_cyto_ oscillations (Figure S1D). In addition, we observed that subthreshold [Ca^2+^]_cyto_ oscillations were preventable by Dan treatment of islets from *db/db* mice. These data further support the role of RyR1 in ER stress-mediated Ca^2+^ dysfunction and show that our results are not limited to findings obtained using chemical stressors solely.

As the selectivity of Dan for blocking RyR1 is controversial (Oo et al. 2015), and RyR2 is the dominant isoform of RyRs in beta cells [certainly by mass (Yamamoto et al. 2019)], we silenced RyR1 and RyR2 individually in INS-1 832/13 cells and found that silencing RyR1 suppressed TM-induced subthreshold [Ca^2+^]_cyto_ oscillations, while comparable silencing of RyR2 did not (Figures 4 and 5). Moreover, silencing RyR1 and RyR2 individually or together failed to prevent UPR activation (Figure 7A-B), in accordance with results we obtained using Dan or ryanodine (Figure 6D-F and Figure 7C-D). These data taken collectively suggest that [Ca^2+^]_ER_ depletion in our system appears to be downstream of UPR activation by TM.

The differential effects of blocking IP3Rs or RyR1 on beta-cell apoptosis as well as SOCE activation and its consequent subthreshold [Ca^2+^]_cyto_ oscillations might be explained by spatial distribution or selective activation, including the potential for dynamic changes, depending on the specific type of ER stress inducer used. The receptors may not be uniformly distributed (Sitia and Meldolesi 1992; Hobman et al. 1998) but selectively and dynamically organized in specific subcellular domains. Stromal interaction molecule 1 (STIM1) and Ca^2+^ release-activated Ca^2+^ channel protein 1 (ORAI1) are the molecular subunits of SOCE channels. STIM1 resides on the ER membrane where it monitors the level of Ca^2+^ in the ER lumen (Stathopulos et al. 2013, 1; Qiu and Lewis 2019, 1). In TM-treated beta cells, RyR1 might be localized nearer to STIM1 than are the IP3Rs, such that STIM1 more readily couples to ORAI1 to mediate Ca^2+^ entry from the extracellular space into the cytosol (SOCE) (40, 41). In contrast, in high glucose-treated beta cells, IP3Rs might also localize closer to STIM1/ORAI1 such that both receptors mediate ER Ca^2+^ depletion and SOCE activation, so that XeC could also inhibit subthreshold Ca^2+^ oscillations, although less efficaciously than Dan (Figure S2D). Resolving the spatial distribution within the ER will require higher resolution imaging approaches that can quantitatively assay the subcellular localizations of RyR1, IP3Rs, STIM1 and ORAI1 under various ER stress conditions.

In summary, the present report demonstrates that RyR1 is a critical player in ER stress-induced ER Ca^2+^ loss and downstream alterations in beta-cell function and viability. Combining these new data with existing knowledge of RyRs and IP3Rs suggests that RyR1 might be a potentially useful therapeutic target for treatment during the onset or progression of type 2 diabetes.

## Materials and Methods

### Materials

Tunicamycin (TM), cyclopiazonic acid (CPA), thapsigargin (TG), caffeine, carbachol and xestospongin C (XeC) were obtained from Cayman Chemical, Ann Arbor, MI. Dantrolene (Dan) was from Sigma-Aldrich, St. Louis, MO. Ryanodine (Ry) was from Abcam, Waltham, MA. RNeasy mini kit for RNA extraction was from Qiagen, Germantown, MD. Superscript RT II was from Invitrogen, Carlsbad, CA. SYBR Green PCR master mix was from Applied Biosystems, Bedford, MA. Primers for qRT-PCR were from Integrated DNA Technologies, Coralville, Iowa. Small interfering RNAs (siRNAs) were from ThermoFisher scientific, Waltham, MA.

### Isolation of pancreatic islets

Pancreatic islets were isolated from male Swiss-Webster mice (3 months of age; 25-35 g) and female *db/db* mice (BKS.Cg-Dock7^m^+/+Lepr^db^/J) and female heterozygous mice at 5-7 weeks of age according to the regulations of the University of Michigan Committee on the Use and Care of Animals, as described (M. Zhang et al. 2003). Islets were cultured in RPMI 1640 medium containing 11 mM glucose, 10% fetal bovine serum (FBS), 10 mM HEPES, 1% penicillin/streptomycin and 1% sodium pyruvate.

### Cell culture and transfection

INS-1 832/13 cells were grown in standard RPMI 1640 medium as described above in 6-well plates at 37°C in a 5% CO_2_ humidified atmosphere and used for experiments after reaching ~70% confluency. INS-1 832/13 cells were transfected with RyR1-specific siRNA, RyR2-specific siRNA or negative control siRNA using lipofectamine RNAiMAX reagent as described in the manufacturer’s protocol (Invitrogen). Transfection was assessed by qPCR and a functional study of RyRs based on Ca^2+^ imaging as described later.

### qPCR

Total RNA was extracted from INS-1 832/13 cells and reverse transcribed to cDNA as described (I. X. Zhang et al. 2020). qPCR was carried out using the primers listed in Table 1, and data were analyzed as described (I. X. Zhang et al. 2020) with expression presented relative to endogenous controls, HPRT1.

**Table 1.**
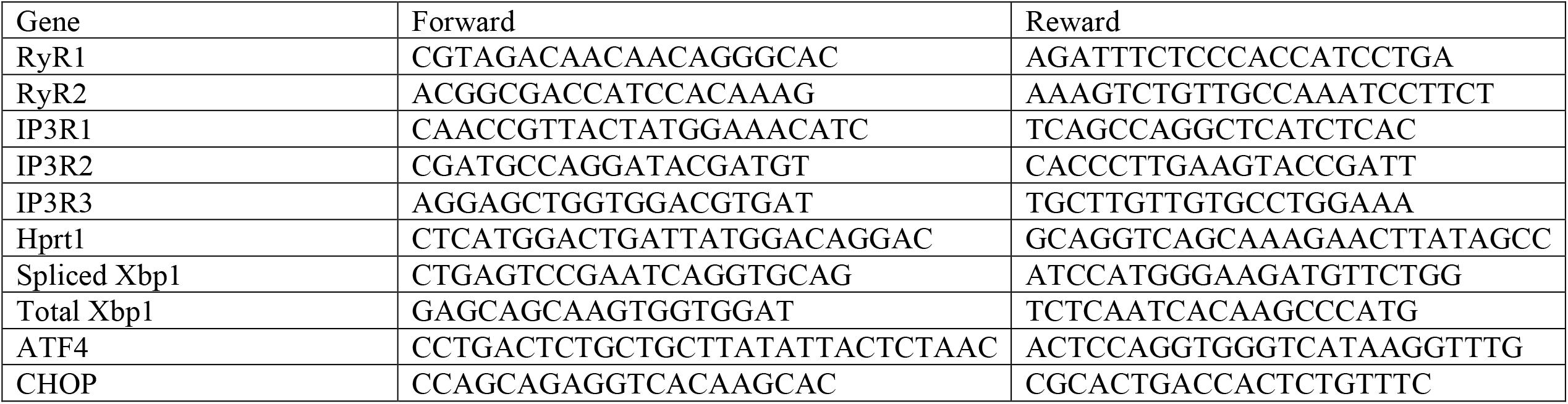
The list of qPCR primer sequences for rat.

### [Ca^2+^]_cyto_ imaging

Islets were loaded with Fura-2/AM (2.5 µM) for 45 minutes in RPMI medium including 5 mM glucose. Islets were then transferred to a 1 ml perfusion chamber containing imaging buffer for 6 min, followed by perfusion at ~1 ml/min. Imaging buffer contained (in mM): 140 NaCl, 3CaCl_2_, 5 KCl, 2 MgCl_2_, 10 HEPES and 5 glucose. Similarly, INS-1 832/13 cells seeded on glass coverslips were loaded with Fura-2/AM (2.5 µM) for 30 minutes in standard RPMI 1640 medium containing 11 mM glucose. Coverslips containing INS-1 832/13 cells were transferred to the perfusion chamber and imaged in imaging buffer. Ratiometric Fura-2/AM imaging was performed using 340/380 nm excitation and collecting 502 nm emission, as previously described (M. Zhang et al. 2003). Fluorescence ratios were acquired using Metafluor software (Molecular Devices, Sunnyvale, CA) and plotted using GraphPad Prism (GraphPad Software, San Diego, CA).

### RyR and IP3R function

Islets or INS-1 832/13 cells were loaded with Fura-2/AM as described above. [Ca^2+^]cyto was measured in imaging buffer without Ca^2+^ for 2 minutes or until the ratio reached a steady state. Cells were then perifused with Ca^2+^free imaging buffer containing either 10 µM caffeine or 100 nM carbachol. Imaging buffer contained (in mM): 140 NaCl, 5 KCl, 3.8 MgCl_2_, 0.1 EGTA, 10 HEPES and 11 glucose.

### [Ca^2+^]_ER_ imaging

[Ca^2+^]_ER_ was measured using an ER-localized FRET biosensor D4ER (Ravier et al. 2011). The same system described above for Fura-2/AM imaging was employed but using 430 nm for excitation and 470/535 nm to obtain ratiometric emission. The imaging solution used contained (in mM): 140 NaCl, 3 CaCl_2_, 5 KCl, 2 MgCl_2_, 10 HEPES, 5 glucose and 0.2 diazoxide (Dz). Dz was included to keep the K_ATP_ channel in its open state to prevent oscillatory Ca^2+^ activity and improve the signal/noise ratio and stability of the ER Ca^2+^ recordings. FRET ratios were acquired using Metafluor software, plotted using Prism, and mean values were calculated using Excel.

### Assays of apoptosis

INS-1 832/13 cells were harvested and prepared for sub-G1 apoptosis assay as described (I. X. Zhang et al. 2020). The percentage of apoptosis was determined by calculating the percentage of cells present in the sub-G1 phase in the DNA content histogram using a flow cytometer housed in the Flow Cytometry Core of the University of Michigan.

### Statistical analysis

Data were expressed as means +/- SEM and analyzed using an unpaired Student’s t-test (Prism) when comparing two groups. Differences between two or more groups were analyzed using two-way ANOVA (Prism) with post hoc multiple comparisons by Tukey’s procedure. Values of p< 0.05 were considered statistically significant.

### Data availability

The data sets corresponding to each figure in the manuscript will be provided as Excel spreadsheets along with details of the statistical methods and statistics for each data set. Excel sheets will be uploaded to the eLife submission site no later than November 10, 2022.

## Acknowledgements

This work was supported by funding from the FastForward program of the University of Michigan Medical School, NIH T32 DK101357 (I.Z.) and NIH RO1 DK46409 (Satin, PI). The authors thank Dr. Arthur Sherman and Dr. Sivakumar Jeyarajan for helpful discussions. The authors declare that they have no conflicts of interest with the contents of this article. I.X.Z., A.H. and J.L. researched data. I.X.Z. wrote the manuscript. A.A. supplied the *db/db* islets. L.S.S. and P.A. reviewed and edited the manuscript.

## Supplemental Figures

**Figure S1.**
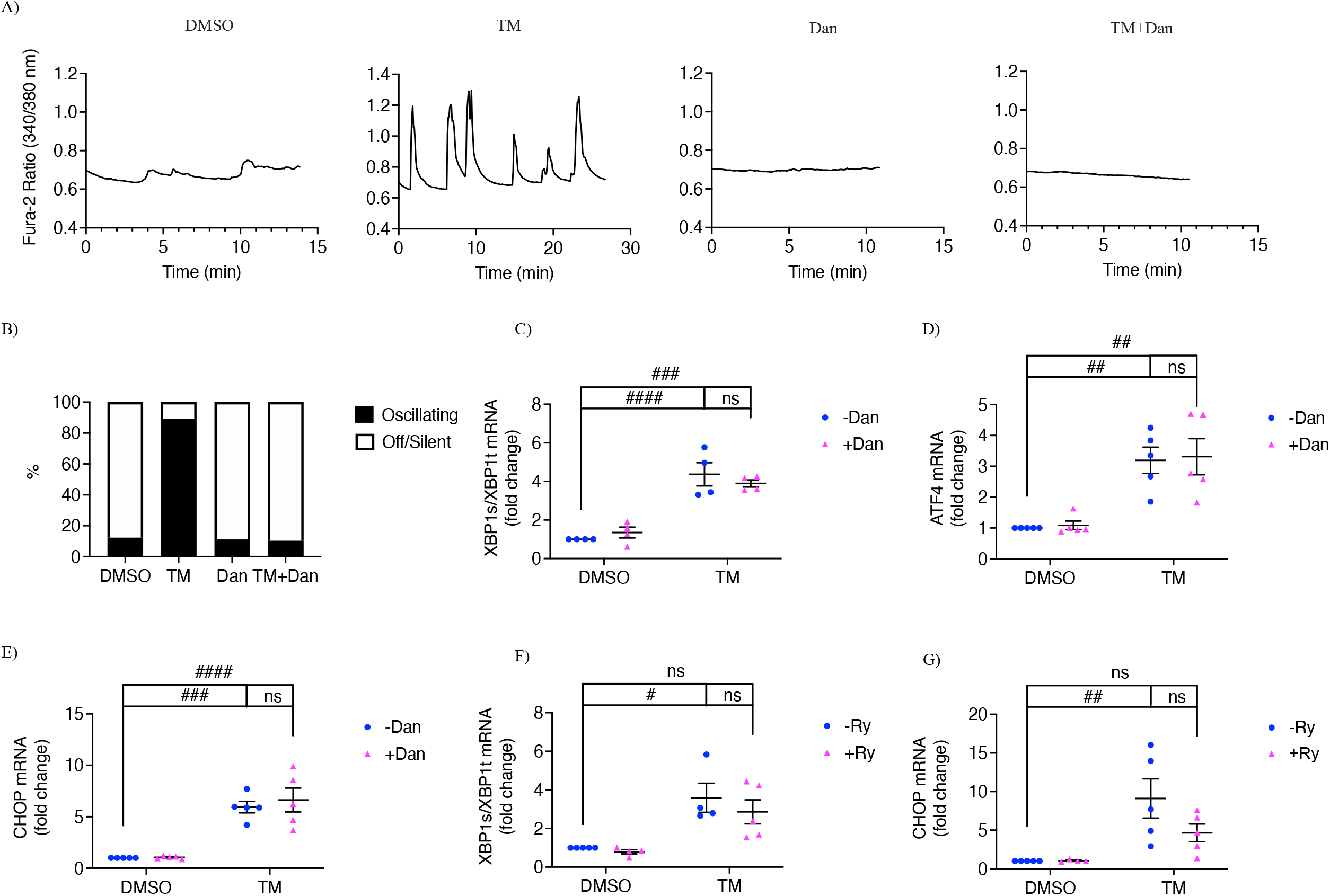
Subthreshold [Ca^2+^]_cyto_ oscillations and UPR activation in beta cells stressed with low dose of tunicamycin. Isolated pancreatic mouse islets were treated with vehicle control (DMSO), tunicamycin (TM, 300 nM), dantrolene (Dan, 10 µM) or TM+Dan for 24 hours in 11 mM glucose islet culture medium. S1A: The responses of [Ca^2+^]_cyto_ to the solution containing 5 mM glucose under the indicated conditions. S1B: Percentage of oscillating islets; n= at least 3 mice. INS-1 832/13 cells were treated with vehicle control (DMSO), tunicamycin (TM, 300 nM), dantrolene (Dan, 10 µM), TM+Dan, ryanodine (Ry, 100 µM) or TM+Ry for 6 hours in INS-1 832/13 culture medium. S1C and S1F: spliced XBP1/total XBP1 ratio; S1D ATF4 and S1E and S1G: CHOP were measured. S1C: Row Factor F(1, 12)= 74.26, p< 0.0001, Column Factor F(1, 12)= 0.03303, p= 0.8588, Interaction F(1, 12)= 1.451, p= 0.2515. S1D: Row Factor F(1, 16)= 36.09, p< 0.0001, Column Factor F(1, 16)= 0.07896, p= 0.7823, Interaction F(1, 16)= 0.001983, p= 0.9650. S1E: Row Factor F(1, 16)= 66.53, p< 0.0001, Column Factor F(1, 16)= 0.3379, p= 0.5691, Interaction F(1, 16)= 0.2465, p= 0.6263. S1F: Row Factor F(1, 18)= 36.33, p< 0.0001, Column Factor F(1, 18)= 1.315, p= 0.2665, Interaction F(1, 18)= 0.6625, p= 0.4298. S1G: Row Factor F(1, 15)= 15.56, p= 0.0013, Column Factor F(1, 15)= 2.201, p= 0.1586, Interaction F(1, 15)= 2.282, p= 0.1517. All values shown are means ± SEM. #, p< 0.05, ##, p< 0.01, ###, p< 0.005; ####, p< 0.0001, ns= not significant; n= at least 4 times repeated per condition, by two-way ANOVA with post hoc multiple comparison by Tukey’s procedure.

**Figure S2.**
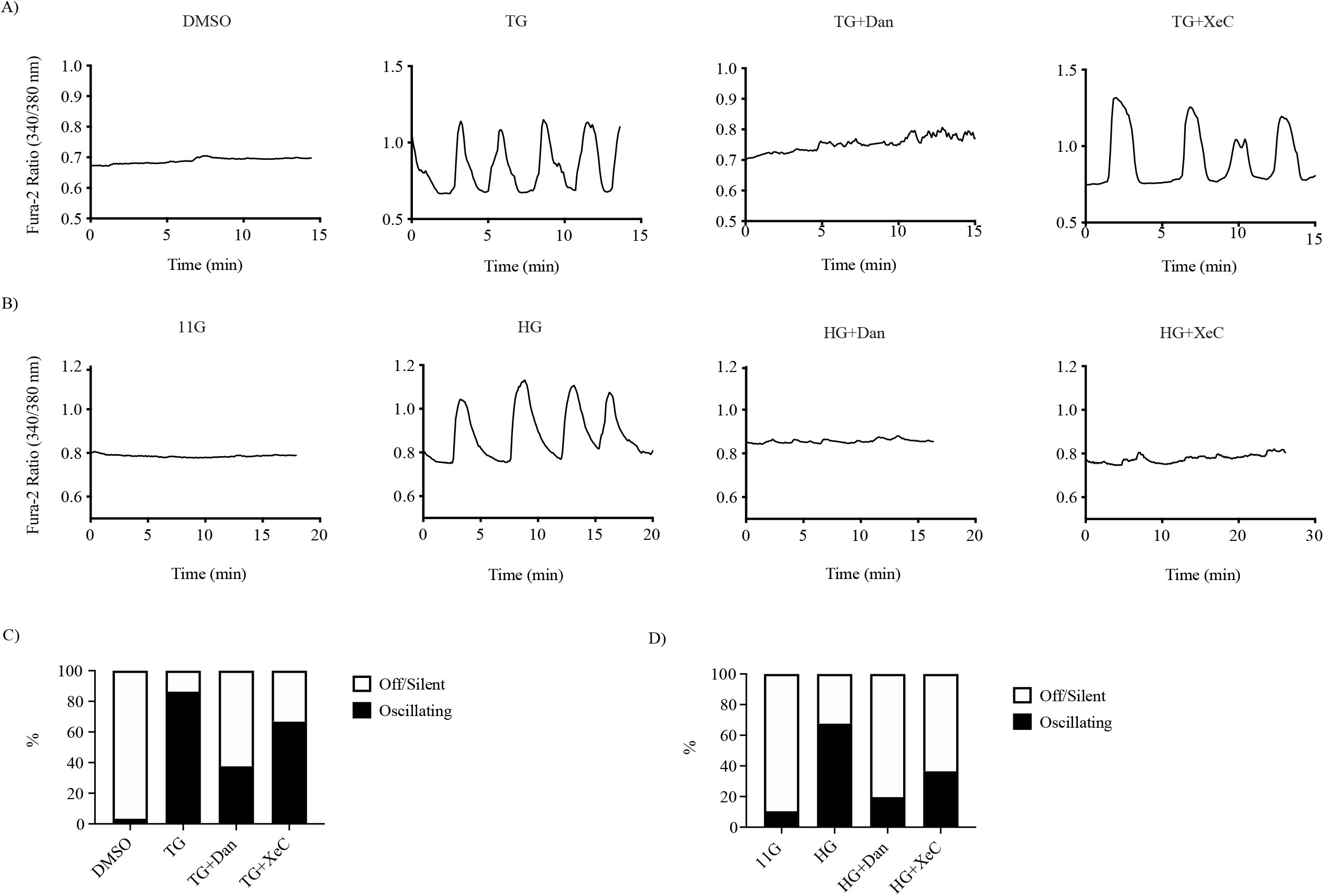
Differential effects of dantrolene and xestospongin C on [Ca^2+^]_cyto_ under subthreshold glucose conditions in islets challenged by alternative ER stress inducers. Isolated pancreatic mouse islets were treated S2A: with vehicle control (DMSO), thapsigargin (TG, 200 nM), TG+dantrolene (Dan, 10 µM) or TG+xestospongin C (XeC, 1 µM); S2B: with control 11 mM glucose, high glucose (HG, 25 mM), HG+dantrolene (Dan, 10 µM) or HG+xestospongin C (XeC, 1 µM) in 11 mM glucose islet culture medium for 16 hours. S2A and S2B: The responses of [Ca^2+^]_cyto_ to solution containing 5 mM glucose under the indicated conditions. S2C and S2D: Percentage of oscillating islets. n= at least 3 mice.

